# Seasonal Changes in Diet and Toxicity in the Climbing Mantella Frog (*Mantella Laevigata*)

**DOI:** 10.1101/361998

**Authors:** Nora A. Moskowitz, Alexandre B. Roland, Eva K. Fischer, Ndimbintsoa Ranaivorazo, Charles Vidoudez, Marianne T. Aguilar, Sophia M. Caldera, Jacqueline Chea, Miruna G. Cristus, Jett P. Crowdis, Bluyé DeMessie, Caroline R. desJardins-Park, Audrey H. Effenberger, Felipe Flores, Michael Giles, Emma Y. He, Nike S. Izmaylov, ChangWon C. Lee, Nicholas A. Pagel, Krystal K. Phu, Leah U. Rosen, Danielle A. Seda, Yong Shen, Santiago Vargas, Andrew W. Murray, Eden Abebe, Sunia A. Trauger, David A. Donoso, Miguel Vences, Lauren A. O’Connell

## Abstract

Poison frogs acquire chemical defenses from the environment for protection against potential predators. These defensive chemicals are lipophilic alkaloid toxins that are sequestered by poison frogs from dietary arthropods and stored in skin glands. Despite decades of research focusing on identifying poison frog toxins, we know relatively little about how environmental variation and subsequent arthropod availability impacts toxicity in poison frogs. We investigated how seasonal environmental variation influences poison frog toxin profiles through changes in the diet of the Climbing Mantella (*Mantella laevigata*). We collected *M. laevigata* females on the Nosy Mangabe island reserve in Madagascar during the wet and dry seasons and tested the hypothesis that seasonal differences in rainfall is associated with changes in the diet and skin toxin profiles of *M. laevigata*. The arthropod diet of each frog was characterized into five groups (i.e. ants, termites, mites, insect larvae, or ‘other’) using visual identification and cytochrome oxidase 1 DNA barcoding. We found that frog diet differed between the wet and dry seasons, where frogs had a more diverse diet in the wet season and consumed a higher percentage of ants in the dry season. To determine if seasonality was associated with variation in frog defensive chemical composition, we used gas chromatography / mass spectrometry to quantify toxins from individual skin samples. Although the assortment of identified toxins was similar across seasons, we detected significant differences in the abundance of certain alkaloids, which we hypothesize reflects seasonal variation in the diet of *M. laevigata*. We suggest that these variations could originate from seasonal changes in either arthropod leaf litter composition or changes in frog behavioral patterns. Although additional studies are needed to understand the consequences of long-term environmental shifts, this work suggests that toxin profiles are relatively robust against short-term environmental perturbations.

## Introduction

Many animals have evolved chemical defenses to deter predators and parasites [1, 2]. Although small molecule chemical defenses are rare among vertebrates [3], some clades of amphibians have evolved alkaloid-based defenses, including Neotropical poison frogs (dendrobatids and bufonids) as well as the less studied clades of mantellid poison frogs from Madagascar, myobatrachids (*Pseudophryne*) frogs from Australia, and eleutherodactylid frogs from Cuba [4, 5]. Indeed, over 800 unique toxins have been identified from poisonous amphibians [6]. Dendrobatid and mantellid poison frogs do not produce alkaloid toxins themselves, but rather sequester them from their diet of leaf litter arthropods [7]. Toxins are stored in specialized skin glands to be actively released in a stress response [5, 8]. Observations from captive breeding colonies of poison frogs led to the dietary hypothesis of poison frog toxicity, as poison frogs raised in captivity lack alkaloid toxins but can acquire them in toxin feeding experiments [9, 10]. As diet and toxicity are tightly linked in poison frogs, there have been many studies attempting to link variation in diet to the striking chemical diversity seen within and among poison frog species [7, 11–15]. Despite progress in understanding these trophic relationships, little is known about how changes in environmental parameters influence the diet and toxicity of poison frogs.

Poison frog sequestration of defensive alkaloid chemicals from arthropods is a remarkable example of convergent evolution that requires innovations in amphibian physiology and behavior. While the best studied clade for chemical defenses are Neotropical dendrobatids [13, 16, 17], some mantellids of Madagascar (genus *Mantella*) also sequester alkaloid compounds from their arthropod diet [9]. Neotropical dendrobatids and Malagasy mantellids shared their last common ancestor nearly 150 million years ago [18] and each are more closely related to undefended clades than to each other. Yet extensive sampling of both lineages over several decades has revealed the presence of hundreds of similar alkaloid toxins [5]. Examples include histrionicotoxins, pumiliotoxins, and decahydroquinolines that target voltage-gated sodium channels and nicotinic acetylcholine channels in the nervous system [19–21]. There is some evidence that toxin auto-resistance has also convergently evolved in dendrobatids and mantellids, where similar mutations in the voltage-gated sodium channel pore have been documented and hypothesized to confer alkaloid resistance in these two frog families [22]. In addition to physiological adaptations, both lineages have evolved behavioral adaptations to support toxicity [23]. Besides diurnal activity patterns, the acquired chemical defenses co-occur with a dietary specialization of ants and mites [13]. Initial reports in some dendrobatid frogs showed ants and mites made up 50-90% of the diet while closely related non-toxic frogs had only 12-16% of their diet consisting of ants and mites [24, 25]. Several species of *Mantella* have a similar abundance of ants and mites in their diet as poisonous dendrobatid frogs [11, 26]. The extent to which toxicity is driven by an active preference for toxic prey [27] and/or simply reflects differences in local arthropod abundance is currently unclear in both clades.

The diversity of arthropod communities in different habitats is hypothesized to explain much of the variation that is seen between different populations of the same poison frog species in both the dendrobatid and mantellid clades [15, 28, 29]. Along with geographic differences in arthropod community composition, seasonal changes in environmental variables also influence arthropod communities. For example, leaf litter moisture plays a large role in leaf litter arthropod distribution and abundance [30–33]. Arthropod species abundance and diversity also drastically change between the wet and dry tropical seasons [34] and across years [33]. Since poison frog toxicity is tightly linked to dietary arthropods, fluctuations in arthropod community composition and abundance likely influence poison frog chemical profiles. Research on the Strawberry poison frog (*Oophaga pumilio*) in Panama has shown there are both geographic and seasonal fluctuations in toxin profiles [16], highlighting the fine spatial and temporal scales at which poison frog chemical defenses can vary. However, the diet contents of these frogs were not reported, and so how variation in toxicity links to variation in diet remains unexplored in the context of seasonal variation.

This study investigated the seasonal fluctuations of diet and toxin composition in the Climbing Mantella (*Mantalla laevigata*), a mantellid poison frog endemic to Madagascar (Fig 1). We tested the general hypothesis that both diet and toxin profiles would change between the wet and dry seasons. To test this hypothesis, we collected *M. laevigata* females on the island reserve of Nosy Mangabe off the northeastern coast of Madagascar in both the wet and dry seasons. We then analyzed stomach contents and quantified toxin profiles of frogs collected in both seasons. Notably, key components of arthropod diet characterization were completed by university freshmen during the ecology module of an experimental science course integrating mathematics, chemistry, biology, and computational approaches into a semester-long lab. Our findings are the first to analyze both diet and toxicity in the context of seasonal variation in poison frogs and contribute to our understanding of how environmental change may impact poison frog diet and chemical defenses.

**Fig 1.**
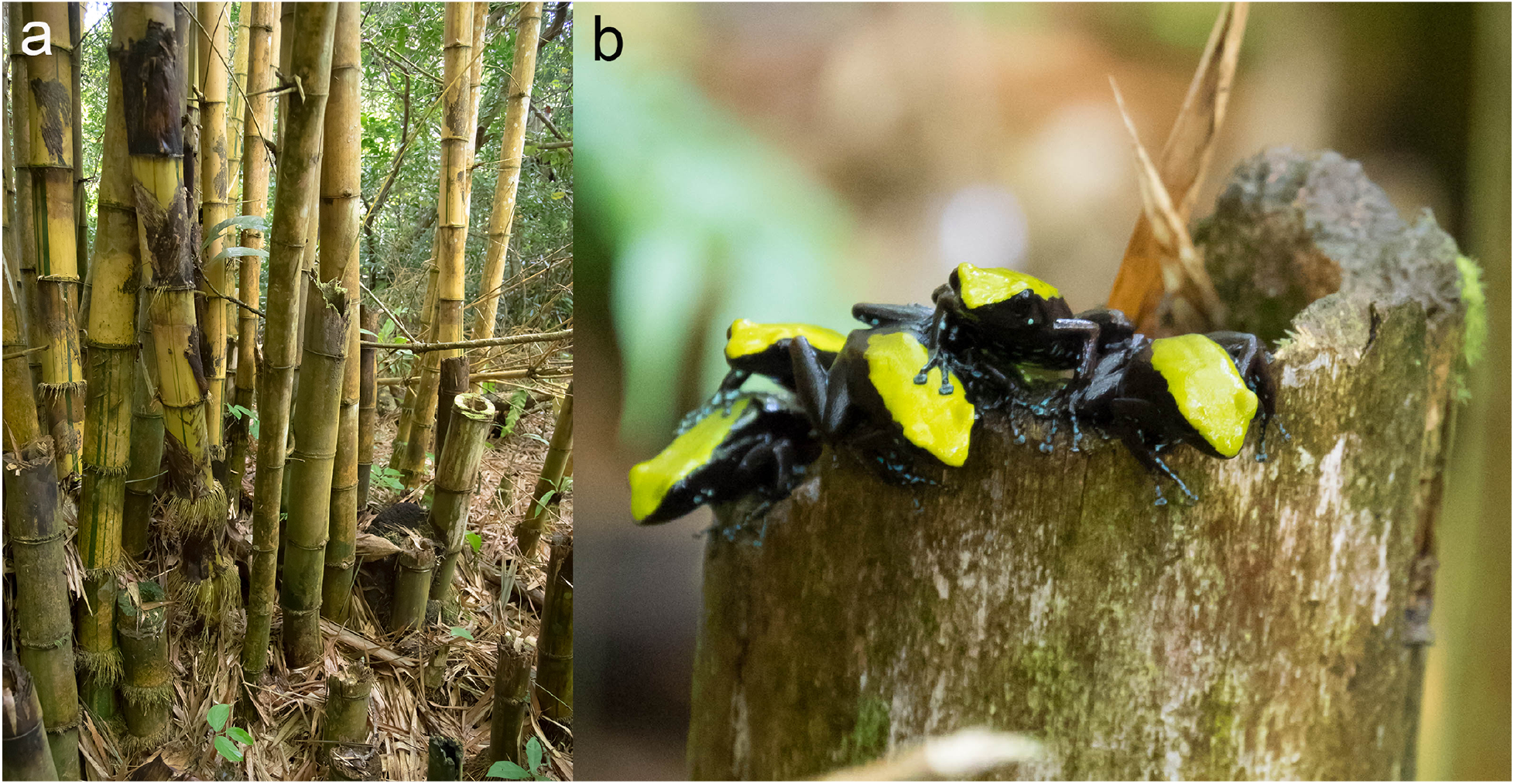
The Climbing Mantella (*Mantella laevigata*) in its habitat. (a) The Nosy Mangabe Reserve contains many areas with large bamboo. **(b)** These bamboo shoots are used by *M. laevigata* as breeding pools.

## Materials and Methods

### Field Collections

Adult female Climbing Mantella (*Mantella laevigata*) were captured on the island of Nosy Mangabe in November-December 2015 during the local dry season (N=11, average mass ± standard deviation: 1.79±0.25g) and July-September 2016 (N=9, average mass ± standard deviation: 1.64±0.17g) during the local rainy (wet) season. Differences in rainfall during these periods reflect local variation in rainfall rather than the major seasons typically associated with Madagascar as a whole (S1 Figure). We collected only adult females for this study because brain tissues were shared with a related study exploring female behavior, allowing us to minimize the number of wild-caught animals collected. Briefly, reproductive females caring for tadpoles were collected in the wet season while non-reproductive females were collected in the dry season. Frogs were collected during the day with a plastic cup and stored individually in plastic bags with air and leaf litter for 25 to 30 min. Individual frogs were photographed, anesthetized with a topical application of 20% benzocaine to the belly, and euthanized by cervical transection. The dorsal skin (from the back of the head but not including the legs) was isolated and stored in glass vials containing 1 ml of 100% methanol. Stomachs were dissected and their contents were stored in 1 ml of 100% ethanol. Remaining frog tissues were preserved in either RNAlater (Life Technologies, Carlsbad, CA, USA) or 100% ethanol. Muscle and skeletal tissue were deposited in the amphibian collection of the Zoology and Animal Biodiversity Department of the University of Antananarivo, Madagascar. Two additional frogs were archived in the Museum of Comparative Zoology at Harvard University (Herpetology A-150719-20). Collection permits (242/15/MEEMF/SG/DGF/DSAP/SCB and 140/16/MEEMF/SG/DGF/DSAP/SCB) were issued by Direction générale des forêts et Direction des aires protégées terrestres (Forestry Branch and Terrestrial Protected Areas Directorate of Madagascar). Exportation of specimens to the United States was completed under CITES permits (1051C-EA12/MG15 and 679C-EA08/MG16). The Institutional Animal Care and Use Committee of Harvard University approved all procedures (protocol 15-03-239).

## Diet Analyses

### Arthropods in Stomach Contents

Arthropod sorting and identification via DNA barcoding was completed by university freshmen during the ecology lab of an integrative science course. Protocols were modified to fit within a three-hour laboratory course and we note all deviations from manufacturer’s instructions below. Stomach contents were placed into a petri dish containing phosphate buffered saline (1X PBS). Individual prey items were sorted and assigned a unique seven-digit identification number with the first four digits being the frog voucher specimen number and the last three digits being the number assigned to the prey item based on the order in which it was isolated. Each arthropod was photographed with a Lumenera Infinity 2 camera mounted on an Olympus dissecting microscope (SZ40) and stored in 100% ethanol for one week. All prey item photos are available on DataDryad (doi: 10.5061/dryad.s7f2gk3).

Diet was quantified by both percent number and percent volume of each arthropod prey type (ants, mites, termites, insect larvae, or “other”) to account for variation in prey size. Volume was determined by taking length and width measurements from photographs imported into ImageJ (National Institute of Health, Bethesda, Maryland, USA). Length was measured from the foremost part of the prey item (including mandibles if applicable) and extended to the rearmost part of the prey item (excluding ovipositors if applicable). Width was measured at the midpoint and excluded appendages. Length and width measurements were then used to calculate the volume of each prey item using the equation for a prolate spheroid: *V* = (4*π*/*3*)*(*Lenghth*/*2*)*(*Width*/*2*)^*2*.

### Molecular Identification of Stomach Contents

DNA from arthropods was isolated using the NucleoSpin Tissue kit (Macherey-Nagel, Bethlehem, PA, USA) according to manufacturer’s instructions with a few deviations to make the protocol amenable to an undergraduate laboratory course. The arthropods were placed in T1 buffer (from the NucleoSpin Tissue kit), crushed with a pestle, and incubated in Proteinase K solution at 56 °C overnight. Samples were then frozen at −20 °C for one week. Extraction of genomic DNA then proceeded according to manufacturer’s instructions and purified genomic DNA was stored at −20 °C for one week. We used PCR to amplify a segment of the cytochrome oxidase 1 (*CO1* or *cox1*) gene from the mitochondrial DNA, the standard marker for DNA barcoding. *CO1* was amplified using the general arthropod primers LCO-1490 (5’-GGTCAACAAATCATAAAGATATTGG) and HCO-2198 (5’-TAAACTTCAGGGTGACCAAAAATCA) from Folmer et al. [35]. For all reactions, we used 2 μL of each primer (10 μM) and 25 μL of 2X Phusion High-Fidelity PCR Master Mix with GC Buffer (New England Biolabs, Ipswich, MA, USA) in a total reaction volume of 50 μL. We used a touchdown PCR program to amplify *CO1* as follows: 95 °C for 5 min; 5 cycles of 95 °C for 30 s, 45 °C for 30 s with −1 °C per cycle, 72 °C for 1 min; and 40 rounds of 95 °C for 30 s, 40 °C for 30 s, and 72 °C for 1 min; ending with a single incubation of 72 °C for 5 min. PCR reactions were stored at −20 °C for one week and then run on a 1% SyberSafe/agarose gel (Life Technologies). Successful reactions with a single band of the expected size were purified with the E.Z.N.A. Cycle Pure Kit (Omega Bio-Tek, Norcross, GA, USA). We successfully amplified and sequenced the target DNA segment from 103 out of 518 (19.9%) of ant samples, 19 out of 82 (23.2%) of mite samples, 1 out of 23 (4.4%) of termite samples, 16 out of 18 (88.9%) of larvae samples, and 27 out of 187 (14.4%) “other” arthropods. Purified PCR products were Sanger sequenced by GeneWiz Inc. (Cambridge, MA, USA). Sequences are available on GenBank (MG947212-MG947379).

We used DNA barcode sequences to qualitatively identify the prey items recovered from stomach contents. Barcode sequences were imported into Geneious (v 11.0.3) for trimming and alignment of forward and reverse sequencing reactions. We used nucleotide BLAST from the NCBI Genbank nr database to identify all arthropods (S2 Table), and nucleotide BLAST from the BOLD Identification System (IDS, http://www.boldsystems.org/index.php/IDS_OpenIdEngine) to identify our *CO1* ant sequences to the species level (S3 Table). This is because the BOLD database is more ant-focused than the Genbank nr database, which increases the likelihood of obtaining a correct match for ant CO1 barcode sequences. We assigned an order, family, genus or species level annotation based on results of the BOLD IDS or Genbank search. We considered results that yielded greater than 96% sequence similarity as sufficient to assign genera or species [36]. For less than 95% BOLD IDS similarity, we assigned specimens to an order or family. For certain specimens, results of the IDS search were more taxonomically ambiguous than others. Some specimens only matched to order, rather than a specific family or genus. For the remaining ants without barcode sequences, photographs were used to identify each whole ant specimen to morphospecies using the barcoded ants as a reference.

## Toxin Analyses

### Isolation of Alkaloids

Alkaloids were extracted as previously described in McGugan et al. [15] and as briefly described below. We extracted alkaloids from individual frog samples, including nine frog skins from the wet season and eleven frog skins from the dry season. Prior to extraction, skin samples were weighed with an analytical scale. The skin and the entire contents of each sample vial (whole tissue piece and methanol) were emptied into a sterilized Dounce homogenizer. To ensure the transfer of all materials, the empty vial was rinsed with 1 ml of methanol and this methanol was also added to the homogenizer. Before homogenization, 25 μg of D3-nicotine (Sigma-Aldrich, St Louis, MO, USA) in methanol was added to each homogenizer to serve as an internal standard. The skin sample was then ground with the piston ten times in the homogenizer before being transferred to a glass vial. The homogenizer was rinsed with an additional 1 ml of methanol in order to collect all alkaloid residues, and this methanol was also added to the final glass vial. All equipment was cleaned with a triple rinse of methanol between samples. Samples were stored at −20°C until further processing.

### Gas Chromatography / Mass Spectrometry (GC/MS)

One ml of each methanol skin extract was evaporated to dryness under nitrogen flow and resuspended in 100 μl of methanol. Samples were analyzed on a Trace 1310 GC coupled with a Q-Exactive Hybrid quadrupole-orbitrap mass spectrometer (ThermoFisher Scientific, Waltham, MA USA). The GC was fitted with a Rxi-5Sil MS column (30 m, 0.25 mm ID, 0.25 μm film, with a 10 m guard column, Restek, Bellefonte, PA, USA). One μl of sample was injected at split 25 in the inlet maintained at 280 °C. The GC program was as follows: 100°C for 2.5 min, then to 280°C at 8°C min^−1^ and held at that final temperature for 5 min. Transfer lines to the MS were kept at 250°C. The MS source was operated in electron ionization (EI) positive ion mode, at 300°C. The MS acquired data at 60000 resolution, with an automatic gain control target of 1×10^6^, automatic injection times and a scan range of 33 to 750 m/z. Mass spectrometry files are available on DataDryad (doi: 10.5061/dryad.s7f2gk3).

## Data Analysis

Except where noted, all analyses were conducted in R, v 3.4.2 using RStudio v1.1.383. All figures were generated using the R package ‘ggplot2’ v 2.21 and aesthetically edited using Adobe Illustrator v 16.0.0. The normality of both the diet and toxin datasets, separated by seasonal groups, were tested with the Shapiro-Wilk test for normality using the R ‘stats’ package v 3.4.2; the fixed effect was consumption and the random effect was the type of prey consumed. Neither set followed a normal distribution. To deal with these deviations from normality, we used a non-parametric Mann-Whitney U tests performed in the R ‘stats’ package v 3.4.2 to compare alkaloid quantities and relative volumes among diet categories between wet and dry season individuals.

### Dietary arthropods

To compare stomach contents across *M. laevigata* populations, we quantified the relative number and volume of all specimens recovered from the stomach contents, including ants, mites, insect larvae, termites and *other* arthropods (S1 Table). Diet variables were not normally distributed and a non-parametric Mann-Whitney U test was used to compare diet categories between wet and dry season individuals. Using the R ‘vegan’ package v 2.5-1, we calculated a Bray-Curtis dissimilarity index for each diet category comparison and plotted the results in two dimensions using a non-metric multidimensional scaling (NMDS). Samples in the resulting NMDS plot were greyscale coded based on season to help visualize the wet and dry seasons frog samples. Polygons were calculated using the ‘ordihull’ function in R using the ‘vegan’ package v. 2.5-1. The purpose of ‘ordihull’ is to create neat, convex outlines to further delineate group separation for visual clarity. A single outline is generated to create a simple polygon that includes all points within an assigned group. We used NMDS twice: to examine the relationship between percent volume and percent number of consumed arthropods between season. We followed our NMDS analyses with permutational multivariate analysis of variance (PERMANOVA) on Bray-Curtis dissimilarities between the samples.

To gain more detailed understanding of what prey the frogs consumed, we performed separate analyses on the three most commonly consumed prey items of ants, arachnids, and *other* categories. For ants, we included in this analisis only whole ants identified to species or morphospecies level by either morphology or molecular barcodes (S3 Table). For arachnids and *other* categories we used prey items that received a reliable CO1 barcode sequence match to the BLASTn database and were classified to order. These data, represented by the percent number of specific prey types eaten within each category (ants, arachnids, and “other”), were not normally distributed. As such, a non-parametric Mann-Whitney U test was used to compare diet categories between wet and dry season individuals.

### Analysis of Toxin Data

Toxins were tentatively identified using the mass spectral data provided in Daly et al [6]. The mass to charge ratio, relative retention times, and relative intensity information was incorporated into a Mass Spectral Transfer File and imported into AMDIS (NIST [37]). This library was used to automatically identify peaks deconvoluted by AMDIS. The identification was weighted by the retention index (calculated from the retention time provided in Daly et al [6], and the retention index of a few easily identifiable compounds like the D3-Nicotine internal standard. Intensities of the model ions for each candidate toxin were extracted from the AMDIS results files and normalized to the mass of skin used for each frog’s alkaloid extraction.

We restricted our analysis to 41 toxins that were present in at least half of the individuals in the study and used this data set to conduct statistical analysis between seasonal groups (S4 Table). The mass spectra of these 41 candidates of interest were manually inspected and compared to the Daly database, as well as to the mass spectra in the NIST14 database, to confirm alkaloid identification. To examine overall alkaloid presence between groups, we calculated a Bray-Curtis dissimilarity index for each alkaloid in the comparison and then plotted the results in two dimensions using a non-metric multidimensional scaling (NMDS). Samples in the resulting NMDS plot were greyscale coded based on season to help visualize how *M. laevigata* individuals differ between the wet and dry seasons in toxin composition. We followed our NMDS analyses with permutational multivariate analysis of variance (PERMANOVA) on Bray-Curtis dissimilarities between the samples. To test if the variance in each group was significantly different, we performed an analysis of similarity (ANOSIM) in the *vegan* R package v 2.5-1. We used a non-parametric Mann-Whitney U test to test for differences in toxin abundance in wet and dry season frogs. We then used permutation testing to control for multiple comparisons among the large number of toxins. Permuted datasets (n=250) were generated by randomly re-assigning wet/dry season labels to individual samples. If the real p-value for a given alkaloid fell in the extreme 5% of the distribution of permuted p-values for that alkaloid, we called that alkaloid differentially abundant between wet and dry season samples.

## Results

### Seasonal differences in diet

We quantified arthropod prey items isolated from *M. laevigata* stomach contents to determine if diet differed between dry and wet seasons (Fig 2). On average, dry season *M. laevigata* individuals consumed 44% more in absolute volume than wet season individuals (S1 Table). We visualized overall differences in diet characteristics using non-metric multidimensional scaling (NMDS) plots (Fig 2a,b). Main prey item classes differed between wet and dry seasons in both percent number (U = 8, *p* = 0.002) and percent volume (U = 6, *p* = 0.001).

**Fig 2.**
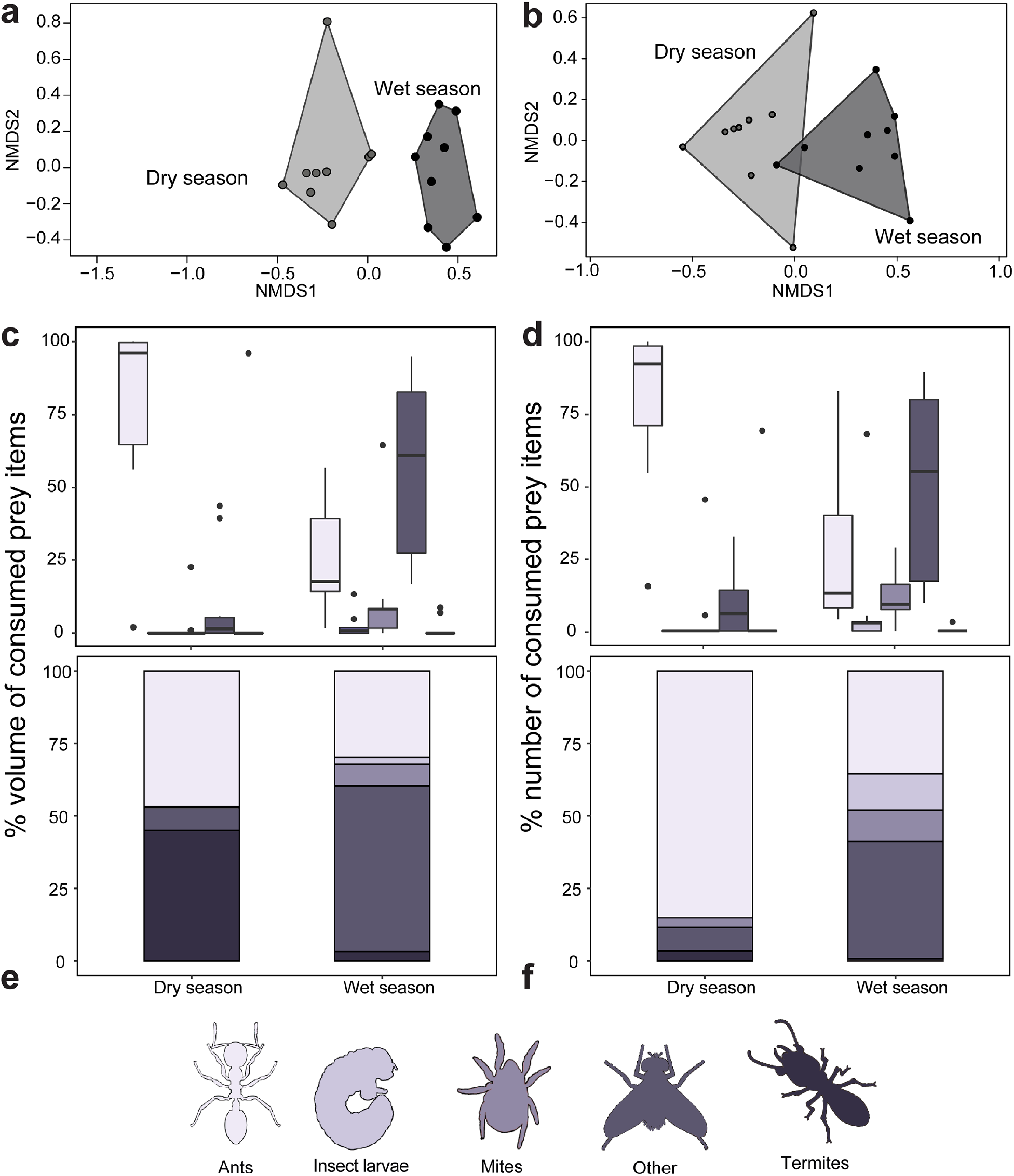
The diet of the Climbing Mantella (*Mantella laevigata*) differs between dry and wet seasons. Non-metric multidimensional scaling (NMDS) biplots based on Bray-Curtis distances display diet differences between seasons using five prey item categories as input (ants, mites, larvae, termites and other prey items) based on **(a)** percent number and **(b)** percent volume of prey items. Below each NMDS plot is abundance in percent volume **(c)** and percent number **(d)** of each prey item category among seasonal groups, and individual distribution of prey item categories among seasonal groups **(e, f)**.

The quantity of ants consumed among wet and dry season groups in *M. laevigata* varied by both relative number (Fig 2c,e, Table 1, S1 Table, U = 90, *p* = 0.002) and relative volume (Fig 2d,f, U = 90*, p* = 0.002). Ants accounted for 84% of the total number of prey items in the dry season, but only 35% in the wet season. Most consumed ants for which we obtained *CO1* sequence are from the subfamily Myrmicinae (95%) and include the tribes Crematogastrini (genera *Cataulacus* and *Tetramorium*), Attini (genera *Pheidole* and *Strumigenys*) and Solenopsidini (genus *Solenopsis*) (Fig 3, S5 Table). In particular, most of the consumed ants that were successfully amplified (90.2%) matched to ants from the genus *Pheidole*. Non-myrmicinae ants were rare but included a *Paratrechina* (99% match to *P. longicornis*, the Longhorn Crazy Ant, Tribe Lasiini) and *Ravavy* MG001, from the subfamily Dolichoderinae (Tribe Tapinomini). Aided by the barcoded ant specimens, we also performed taxonomic identification of all whole ants by morphology (S3 Table). The three most common species identified were *Pheidole* sp. MG051, *Pheidole* sp. MGS128, and *Pheidole* nr. *madecassa* (Fig. 3). Between seasons, consumption of these three ant species did not differ (*Pheidole* sp. MG051; U = 36, *p* = 0.7939, *Pheidole* sp. MGS128; U = 42, *p* = 0.199, *Pheidole* nr. *madecassa*; U = 42, *p* = 0.199).

**Table 1.** Broad diet characterization of wet and dry season groups of *Mantella laevigata*.

|  | % number |  |  |  |  | % volume |  |  |  |  |
| --- | --- | --- | --- | --- | --- | --- | --- | --- | --- | --- |
|  | <u>ants</u> | <u>mites</u> | <u>larvae</u> | <u>termites</u> | <u>other</u> | <u>ants</u> | <u>mites</u> | <u>larvae</u> | <u>termites</u> | <u>other</u> |
| **Dry** | 85.1 | 3.4 | 0 | 3.4 | 8.1 | 46.9 | 0.5 | 0 | 45.0 | 7.6 |
| **Wet** | 35.5 | 10.8 | 12.5 | 0.8 | 40.3 | 29.9 | 7.4 | 24 | 3.2 | 57.2 |
| **U** | 90 | 15.5 | 16.5 | 44 | 9 | 90 | 13.5 | 16.5 | 44 | 6 |
| **p** | 0.002 | 0.007 | 0.002 | 0.541 | 0.002 | 0.002 | 0.004 | 0.002 | 0.541 | 0.001 |

**Fig 3.**
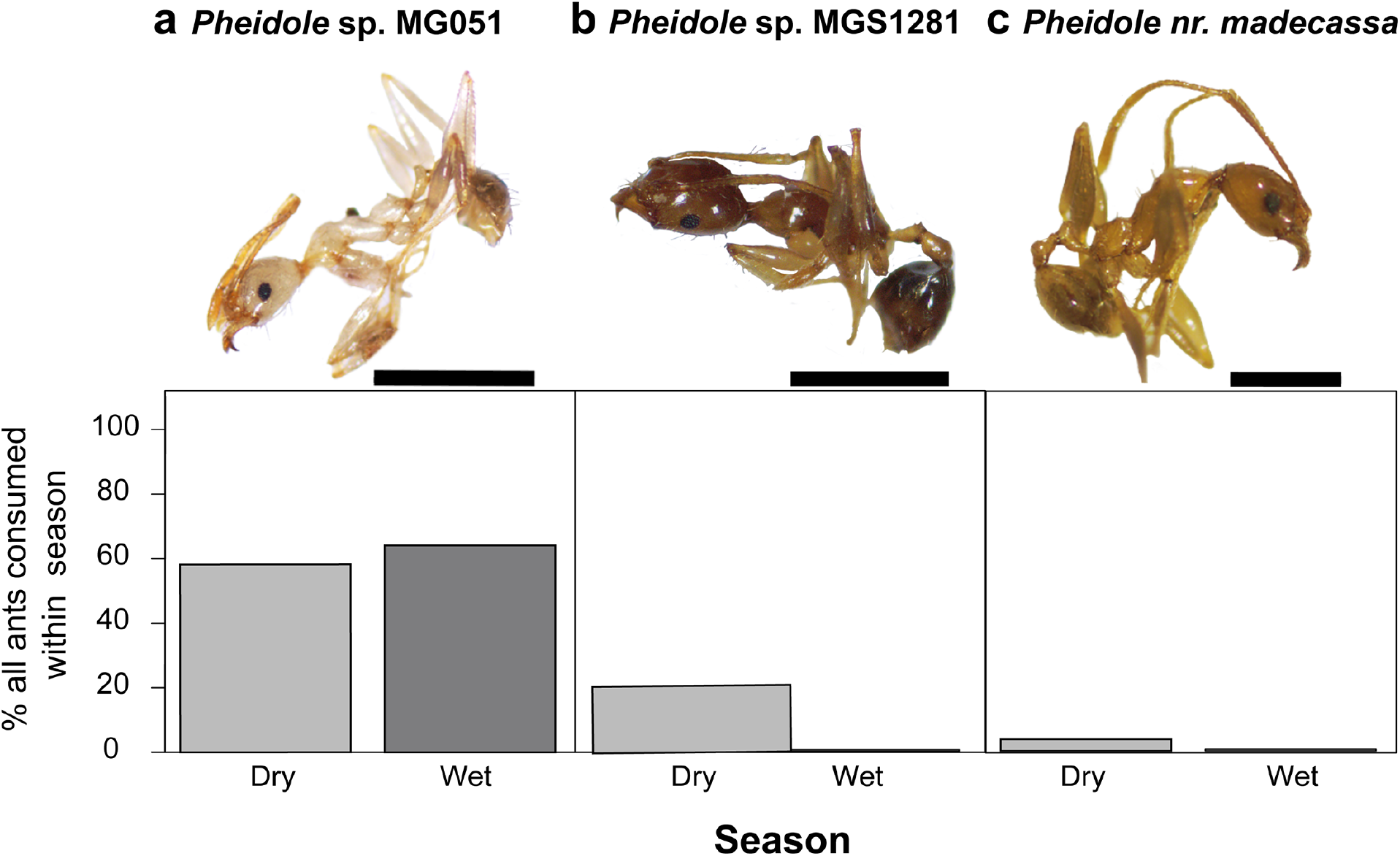
Identification and abundance of the three most common ant prey species of *Mantella laevigata*. Relative percentages of the three most commonly consumed, morphologically identifiable ant species does not differ between wet season and dry season frogs (Mann-Whitney U). Scale bar = 1.0mm, for all individuals. **(a)** *Pheidole* sp. MG051 **(b)** *Pheidole* sp. MGS128, **(c)** *Pheidole* nr. *madecassa.*

Frogs collected in the wet season consumed more mites in both relative volume (U = 13.5, *p* = 0.004) and relative number (U = 15.5, *p* = 0.007) compared to frogs collected in the dry season (Fig 1c-f, Table 1). Barcode information for arachnids is limited compared to ants and so classifications were restricted to Order for all specimens. Most of the consumed arachnids for which we obtained *CO1* sequence are mites (Subclass Acari) from the Sarcoptiformes order (94.7%) and a single mite from the Mesostigmata order (Fig 4, S6 Table). Several spiders were also recovered from the stomach contents of wet season frogs. Consumption of the three most consumed, genetically identifiable arachnid orders did not differ between seasons (Sarcoptiformes; U = 4, *p* = 0.5338, Mesostigmata; U = 2, *p* = 0.999, Araneae; U = 1.5, *p* = 0.729).

**Fig 4.**
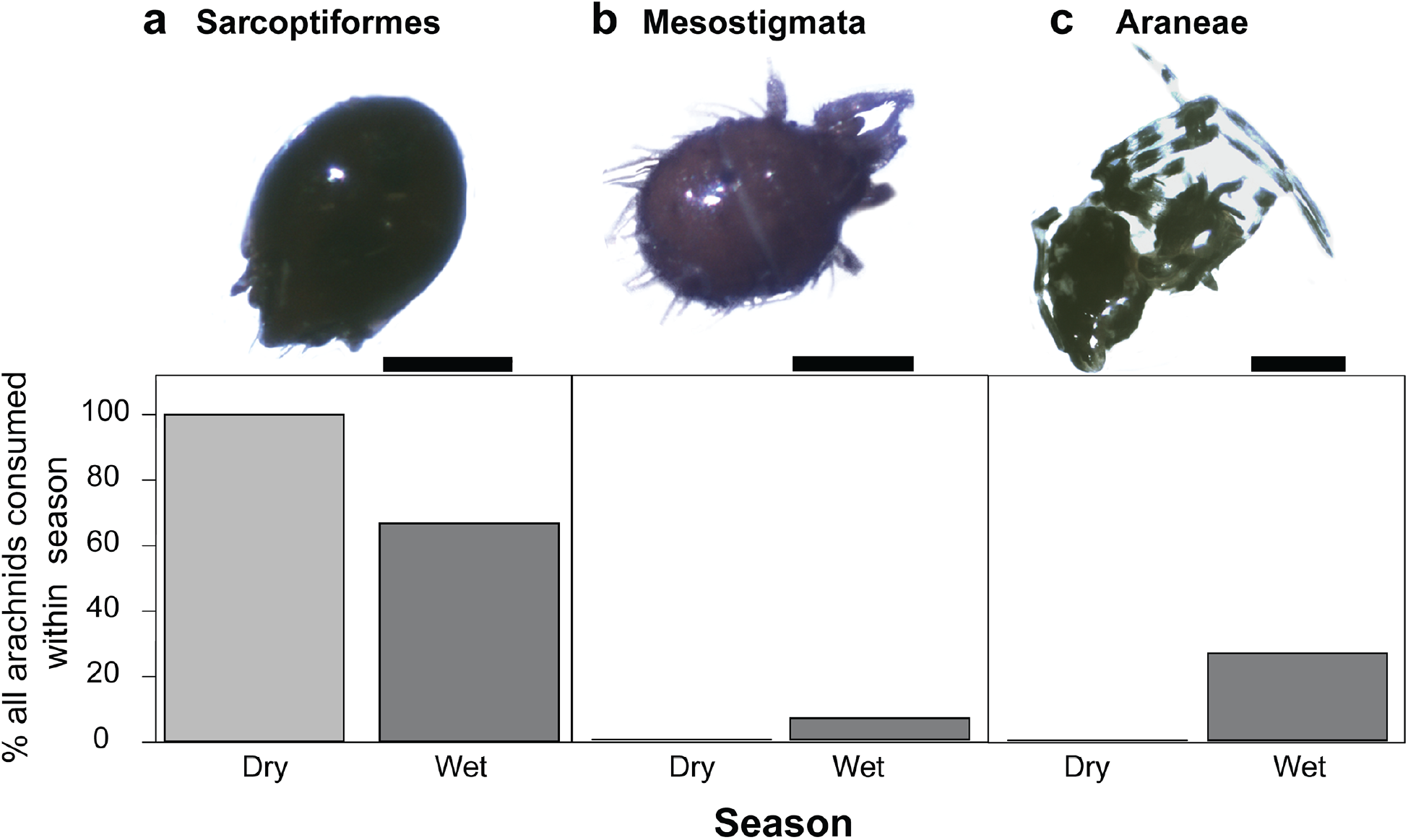
Identification and abundance of the three most common arachnid prey species of *Mantella laevigata*. Relative percentages of the three most commonly consumed, genetically identifiable arachnid orders do not differ between wet season and dry season frogs (Mann-Whitney U). **(a)** Sarcoptiformes; scale bar = 0.5mm. **(b)** Mesostigmata; scale bar = 0.25mm. **(c)** Araneae; scale bar = 0.25mm.

Insect larvae made up a substantial portion of the stomach contents of frogs collected in the wet season but constituted no part of dry season frog diet (Fig 1a-e; Table 1; S7 Table; volume: U = 16.5, *p* = 0.002; number: U = 16.5, *p* = 0.002). Most of these insect larvae were flies (45%, Order Diptera), including many midge larvae (Family Ceratopogonidae), although some wood moth larvae (Order Lepidoptera, Family Crambidae) were also present. We grouped all other arthropods recovered from the stomach contents into an “other” category, in which we noted the presence of several beetles (Order Coleoptera), including a number of click beetles (Family Elateridae). Other arthropods included springtails (Collembola), parasitic wasps (Order Hymenoptera, Family Braconidae), termites, and barkflies (Order Psocoptera). Frogs collected in the wet season consumed more of these “other arthropods” in both volume (U = 6, *p* = 0.001) and number (U = 9, *p* = 0.002) compared to frogs collected in the dry season (Fig 1c-f, Table 1). There were no significant differences in the abundance of the three most consumed, genetically identifiable “other” orders between wet and dry season frogs (Fig 5; Collembola; U = 7, *p* = 0.8026, Coleoptera; U = 13, *p* = 0.2284, Diptera; U = 8, *p* = 0.1).

**Fig 5.**
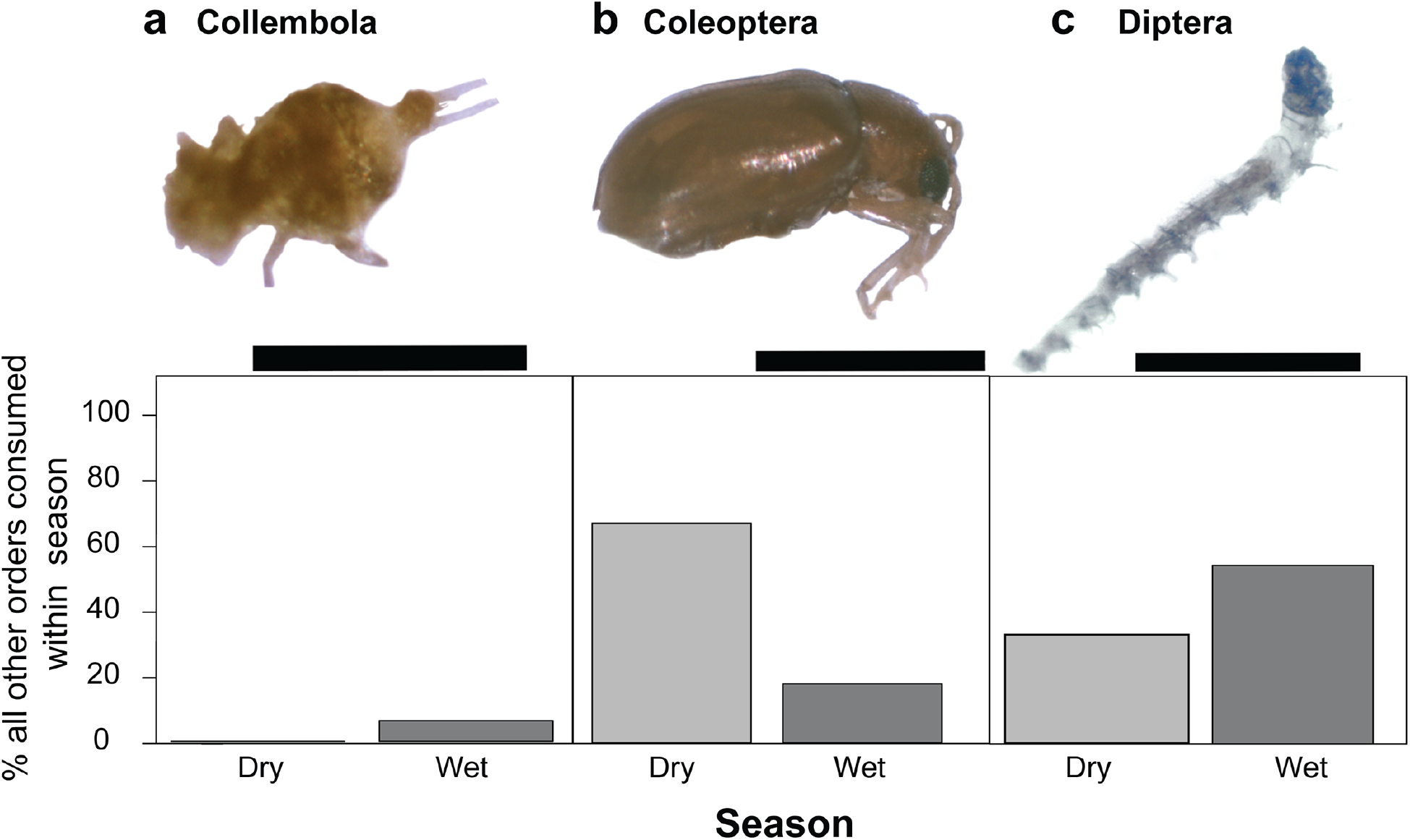
Identification and abundance of the three most common other prey species of *Mantella laevigata*. Relative percentages of the three most commonly consumed, genetically identifiable other (non-ant or -arachnid) prey items does not differ between wet season and dry season frogs (Mann-Whitney U). **(a)** Collembola; scale bar = 0.5mm **(b)** Coleoptera; Scale bar = 1.0mm **(c)** Diptera; scale bar = 1.0mm

### Seasonal differences in toxin profiles

We identified 41 alkaloids in *M. laevigata* that were present in at least half of the frogs included in this study (S4 Table). Several alkaloids are likely ant-derived [38] (S8 Table), including the following eight decahydroquinolines (DHQ), one tricyclic alkaloid and one 3,5 di-substituted pyrrolizidine (3,5-P): **189** DHQ,**193D** DHQ, **195A** DHQ, **195J** DHQ, **209A** DHQ, **211A** DHQ, **211K** DHQ, **219A** DHQ, **237O** Tricyclic, **239K** 3,5-P. Others are likely mite-derived [38] (S8 Table), including pumiliotoxins (PTX): **197G** 5,8,6-I (5,6,8-trisubstituted indolizidine), **211L** 5,8,6-I, **225K** 5,8,6-I, **239W** 5,8,6-I, **267W** 5,8,6-I, **253F** PTX, **275E** 5,8,6-I 2^nd^ Isomer, **291E** DeoxyPTX 1^st^ Isomer, **305A** aPTX (allopumiliotoxin), **307A** PTX pumiliotoxin A minor isomer, **307G** PTX, **321B** hPTX (homopumiliotoxin), **321D** hPTX 1stIsomer, **323A** PTX pumiliotoxin B, **337B** hPTX.

Frogs collected in different seasons had no overall differences in toxin profiles (NMDS analysis, F=1.279; p = 0.22; Fig 6a). Overall wet season frogs appear less variable in their alkaloid profiles, clustering as if their toxin profiles are a subset of dry season toxins, but this variation between seasonal groups was not statistically significant (R = 0.03497; p = 0.266). However, seven individual toxins differed significantly in abundance between wet and dry season groups (Mann-Whitney U tests, Figure 5b): **211A** DHQ, (U = 16.5, *p* = 0.01), **211K DHQ, (U = 12, *p* = 0.004), 211L** 5,8,6-I (U = 11, *p* = 0.003), **253F** PTX (U = 13, *p* = 0.004), **275E** 5,6,8-I (U= 17, *p* = 0.01), **305A** aPTX (U = 80, *p* = 0.02), **323A** PTX (U = 9, *p* = 0.001). Six of these toxins were higher in abundance in wet season frogs compared to dry season frogs.

**Fig 6.**
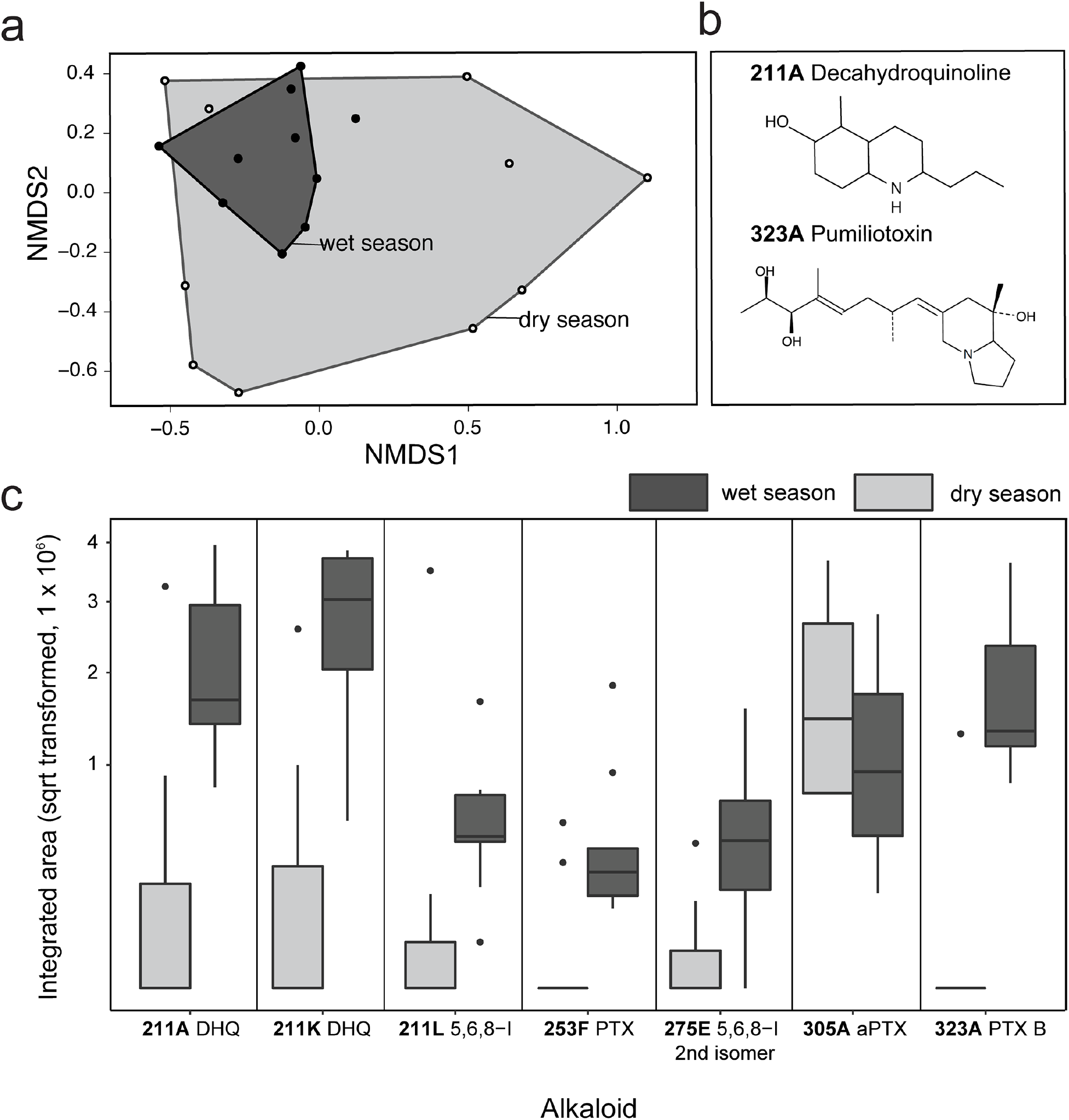
Seasonal comparison of chemical defenses in *Mantella laevigata*. (a) Non-metric multidimensional scaling (NMDS) biplots based on Bray-Curtis distances show overlap of toxin profiles between wet and dry season frogs. **(b)** Chemical structures of two toxins that differ in abundance between collected in the wet and dry season. **(c)** Abundance of seven alkaloids that are significantly different in *M. laevigata* frogs collected in the dry and wet seasons. Box plots show the median, first and third quartiles, whiskers (±1.5 interquartile range) and outliers (black dots). PTX = Pumiliotoxin; DHQ = Decahydroquinoline; 5,6,8-I = 5,6,8-trisubstituted indolizidine.

## Discussion

We showed that arthropod consumption differed between groups of *M. laevigata* from the wet and dry seasons both within and across prey types (ants, mites, larvae, and other). Although there were no overall differences in the number and types of toxins present between seasons, seven unique alkaloids differed in abundance between seasons with the majority more abundant in the wet season. We discuss our diet and toxin results in the context of studies in other poison frogs and link these two measurements together in the context of trophic ecology and environmental change.

### Diet of the Climbing Mantella compared to other poison frogs

The diet of poison frogs has been intensely studied since it was demonstrated that poison frogs acquire their toxins from their diet. Our study is the first in mantellids to compare diet across wet and dry seasons and found that frogs in the wet season have a broader diet that includes an abundance of fly larvae, while frogs in the dry season eat more ants. Other studies have examined the diet and food preferences of *Mantella* poison frogs and have found variation between study sites and species. Two studies across numerous *Mantella* species (*M. baroni, bernhardi, betsileo, haraldmeieri*, *laevigata*, *madagascariensis, nigricans*) showed ants comprised an average of 27-91% of prey volume in the wet season [26, 39]. In contrast, a study with the Golden Mantella (*M. aurantiaca*) showed ants made up a low percentage of the diet (11-15%) and mites a higher percentage (18-34%) in the wet season [27]. Together with a food preference assay showing that *M. laevigata* will readily eat any small prey items available in the leaf litter (rather than having a continuous and active preference for ants and mites) [26], these observations suggest that some mantellids – including our focal species – are not ant-specialists. Despite the overall lower percentage of ants in the diet of *M. aurantiaca* and *M. laevigata* in the wet season, both species showed high selectivity for *Pheidole* ants among the ants they did consume [27]. Specifically, over half of the consumed ants in both the wet and dry seasons were a single ant species, *Pheidole* sp. MG051 (Fig 3). This suggests *M. laevigata* frogs may prefer certain types of ants despite not specializing on this prey type. Alternatively, there may be more *Pheidole* ants present in the leaf litter and the frogs are sampling the most abundant species. Detailed leaf litter and prey preference assays are needed to distinguish between these alternative explanations. In summary, our observations in *M. laevigata* are consistent with previous studies in *M. laevigata* and *M. aurantiaca,* and demonstrate that some *M. laevigata* are active foragers that will readily eat nutritious prey items like flies or insect larvae. They appear to retain a preference for certain types of ants over others, although testing this hypothesis in future studies would require data on prey availability.

Ecological studies in tropical habitats have shown that arthropod diversity and abundance change in the leaf litter based on moisture [31]. Although we did not directly survey leaf litter arthropods in this study, we suggest that seasonal variation in rainfall influences the leaf litter arthropods available for frogs to prey on. While many studies have reported that Neotropical poison frogs in the Dendrobatidae family are selective for the ants and mites that typically make up 80-90% of their diet [13, 25], the relative proportion of these prey categories can vary by season. For example, in the Strawberry poison frog *Oophaga pumilio*, the relative proportion of ants and mites varied from 30% ants / 65% mites in summer months to 75% ants / 10% mites in winter months [25]. A study focusing specifically on seasonal variation in poison frog diet found that French Guianese *Dendrobates tinctorius* had a more diverse arthropod diet in the wet season compared to the dry season [40]. A study focusing on *M. aurantiaca* supports this idea in mantellids [27] where there were a large number of flies (Diptera) and springtails (Collembola) among consumed prey items in the wet season. Our findings that *M. laevigata* consumed a greater diversity of arthropods in the wet season align with these studies and support the idea that increased arthropod diversity in the wet season leads to increased diet diversity in both dendrobatids and mantellids. In contrast to *M. laevigata*, however, *O. pumilio* and *D. tinctorius* still had an ant-heavy diet in both the wet and dry seasons lending further support to the assertion that *M. laevigata* are not strict ant specialists.

In addition to variation in prey availability, seasonal changes in diet may also result from seasonal behavioral differences. Behavior may change seasonally based on the microhabitats frogs inhabit as humidity and temperature change. In our study, we found that dry season *M. laevigata* frogs consumed 44% more prey than wet season frogs (S1 Table). During the warmer dry season, we observed more frogs foraging in places that were more shaded, and likely cooler, such as under water pipes and decaying logs at our field sites, and these areas may also harbor more arthropods. Thus, seasonal diet changes may result from changes in prey availability in conjunction with behavioral changes.

### Defensive Chemicals in the Climbing Mantella

We reported 41 alkaloid toxins that were present in at least 50% of the individuals in this study. Many toxins overlap with those that have been reported previously in mantellids and other Neotropical poison frogs (S4 Table). Ten alkaloids have never been reported in a *Mantella* species to our knowledge [28, 39, 41], and include one unclassified alkaloid (**153B**), three decahydroquinolines (**209A**, **211A**, **211K**), a 5,6,8-trisubstituted indolizidine (**211L**), a 5,8-disubstituted indolizidine (**225D**), an izidine (**233B**), 3,5-disubstituted indolizidines (**237E**), a tricyclic (**237O**), and a quinoline (**380**). Most of these alkaloids had previously only been found in dendrobatid poison frogs, many of which had been reported in Panamanian *O. pumilio* [16, 29]. The tricyclic **237O** alkaloid had previously been reported in a bufonid [42]. Our findings add support to previous reports of convergent evolution of alkaloid sequestration in Malagasy and Neotropical poison frogs [11].

Although we found significant seasonal variation in the diet of *M. laevigata*, we did not find drastic seasonal variation in toxin profiles in terms of the number and quantity of toxins present. There is a trend for wet season frogs to have less variable toxin profiles than dry season frogs, but this variance is not statistically significant (Fig 6). These observations are in contrast to another study that analyzed seasonal variation in alkaloid toxins in dendrobatid poison frogs, where Saporito et al [16] found seasonal variation in the absence/presence of toxins in Panamanian *O. pumilio*. However, the *O. pumilio* study included many collection sites and individuals whereas our study was focused on a single population of *M. laevigata* and had a smaller sample size. Thus, seasonal variation in concert with geographical variation may be contributing to greater differences in *O. pumilio* than those we observed in *M. laevigata*. Moreover, adult poison frogs can retain their toxins for months and even years in captivity [43], and the ability to retain toxins for long periods of time could also explain the buffering effect we see in the present study, where frog toxin profiles look similar across seasons despite changes in diet. Based on the changes in abundance of specific toxins, we hypothesize that seasonal changes in diet do indeed influence toxicity (i.e toxin abundance), but that overall toxin profiles (i.e. toxin presence/absence) are stable due to long-term toxin retention across seasons. Research involving more frequent tracking of overall toxicity in frogs is needed in order to gain a deeper understanding of alkaloid toxin retention and the temporal relationship between toxin consumption, storage, and potency, especially in the context of natural seasonal variation.

### Connecting diet to toxicity in the context of seasonal variation

Testing the association between alkaloid diversity and stomach contents is difficult due to the limited sampling of frogs in this study. Here we remark on our observations between prey items recovered from the stomach contents and published alkaloid profiling in these arthropods. The bulk of toxins found in *M. laevigata* from the present study originate from myrmicine ants and have previously been found in both dendrobatids and mantellids [6]. Many decahydroquinolines were found in both seasonal groups, most of which are known to originate from myrmicine ants [12, 14]. Myrmicine ants from the genus *Pheidole* account for the majority of ants that *M. laevigata* consume across seasons and this suggests that *M. laevigata* may rely on ants for their chemical defenses. Of the seven toxins that were significantly different in abundance between seasons, three have a proposed ant origin (S8 Table). Several ants recovered from the *M. laevigata* diet have been documented to contain toxins, including *Tetramorium* ants that carry pumiliotoxins and 5,8 indolizidines [11] and *Paratrechina* ants that carry a variety of alkaloids [44]. Some *Solenopsis* specimens were also recovered from the stomach contents, and these ants have been well-studied in the context of fire ant defensive alkaloids [45]. Both the number and volume of ants consumed by *M. laevigata* varied between seasons and this could explain some of the seasonal variation in toxin abundance. Given most of the ants recovered from the stomach contents were from the genus *Pheidole*, a more systematic characterization of the chemical composition of these ants could help to confirm this ant genus as a dietary source of alkaloid toxins in poison frogs. Such a study could also reveal dietary preferences at the morphospecies level, as have been reported in a previous study of *M. aurantiaca* [27], which could be correlated with the acquisition of specific toxins. Strong selectivity for *Pheidole* ants even in the absence of an overall dietary specialization on ants may allow for simultaneous diet diversification and toxin maintenance.

Mites are known to be a rich dietary source of alkaloids in poison frogs [46]. At least 80 alkaloids have been documented from extracts of oribatid mites [15, 46, 47], corresponding to 11 of the 24 classes of alkaloids found in dendrobatid poison frogs. We found three families of oribatid mites known to contain alkaloids in *M. laevigata* stomach contents: Galumnidae, Scheloribatidae, and Oppiidae. Tricyclic alkaloids have been reported in Galumnidae mites while pumiliotoxins have been reported in both Schelorbatidae and Oppiidae mite families [46, 48]. Thus, although mites made up a small proportion of the *M. laevigata* diet, they are likely an important source of toxins and it is notable that their number and volume in the stomach contents differed across seasons. Out of the seven toxins that significantly varied in abundance in *M. laevigata* across seasons, three are thought to have a mite origin (5,6,8-trisubstituted indolizidine **275E**, allopumiliotoxin **305A**, and pumilitotoxin B **323A,** see S8 Table) [6]. Compared to ants, little is known about the taxonomy and chemistry of mites and more work is needed on this group of organisms to better understand the trophic chain of toxin sequestration.

Although many toxin classes have a proposed ant or mite origin [6], many alkaloid toxins have not yet been documented in specific leaf litter arthropods and generally very little is known about which specific species of ants, mites, or other arthropods carry the alkaloids found in poison frogs. Of the 41 toxins documented in *M. laevigata* in the present study, 20 have an unknown origin. Further chemical analyses are needed to trace the origin of these toxins to specific arthropod species in the trophic chain. In addition, analyses of prey items recovered from stomach contents represent only a snapshot in time and long-term monitoring of diet is needed in order to make conclusions about alkaloid origins in arthropods. Future studies that combine long-term diet monitoring with behavioral testing and chemical analyses of arthropods from the leaf litter will shed light on how prey abundance interacts with prey preference to drive variation in poison frog toxin profiles.

### Conclusions

We found substantial variation in the *M. laevigata* diet between seasons. Although we used both molecular and morphological methods to assay diet in the present study, these results are only a snapshot in time and repeated sampling of the same individuals by stomach flushing or fecal barcoding would be a valuable step forward in understanding diet variation a finer scale. Moreover, leaf litter studies coupled with diet analyses would be valuable in assessing the intersection of prey preference, prey availability, and frog behavior. Finally, although there were differences in the abundance of specific toxins between frogs collected in the wet and dry season, overall toxin profiles were similar across seasons. The ability for adult poison frogs to retain their toxins for years in captivity coupled with our observations of cross seasonal stability in toxin profiles suggests that poison frog defenses are buffered against acute environmental changes. Nonetheless, changes in toxin abundance even at the short timescale we consider here suggest that more extreme, long-term shifts in temperature and humidity will impact poison frog toxicity and continued long-term monitoring of poison frog populations is warranted. Given the unique and fascinating link between arthropod diversity and poison frog biology, poison frogs may serve as a useful model for understanding the ecological consequences of global environmental change across trophic levels.

## Acknowledgements

We would like to thank the University of Antananarivo, ICTE/MICET, and Madagascar National Park (MNP) for providing logistical assistance. We thank Hadley Weiss for technical assistance. We are very grateful to the Malagasy authorities for research permits. We would like to thank the local MNP guides Béonique, Donné, Clotric, and Marcel for their help and company.

## Author Contributions

LAO and MV designed the research; ABR, EKF, and NR collected samples in Madagascar; freshman Harvard College students (LS50:Integrated Science; MTA, SMC, JC, MC, JPC, BD, AHE, FF, MG, EYH, NSI, CL, BL, NAP, KKP, LUR, DAS, YS, and SV funded by the Howard Hughes Medical Institute Professor’s Award 520008146 to AWM) performed the molecular arthropod identification under the guidance of EKF, ABR, and LAO as part of a course laboratory module on trophic ecology; EA and NAM also performed molecular arthropod identification; DAD identified ant specimens to species/morphospecies; NAM performed the extraction of alkaloids from skin samples; CV and SAT performed the GC/MS experiments; NAM and CV analyzed the GC/MS data; NAM analyzed the diet data; NAM and LAO wrote the paper with contributions from all authors.

## Supporting Information

**S1 Figure. Environmental information for Sambava, Madagascar.** (a) Sambava is roughly 150 km northeast of Nosy Mangabe. Precipitation (blue) and temperature (orange) are shown for 2015 **(b)** and 2016 **(c)**; field work time is indicated in red. Environmental data was obtained from: https://www.historique-meteo.net/afrique/madagascar/sambava/

**S1 Table. Diet data for each *Mantella laevigata* frog.** The absolute and relative volume and number for each prey category for each frog is listed and summarized.

**S2 Table. Arthropod sample barcode information.** The details for each cytochrome oxidase 1 barcoded non-ant arthropod is listed with the sample ID, the prey type, and information on the top BLAST hit including the species name, percent identity, and E value.

**S3 Table. Morphological and genetic identification of ant specimens.** For all morphologically or genetically identifiable ant specimens found in *M. laevigata* stomach contents across wet and dry seasons, we list the sample ID, species name, BOLD and Genbank database IDs and percent matches to these IDs For Genbank-matched individuals, BLASTn hit, specimen Genbank accession number, percent identity and E value are also listed.

**S4 Table. Abundance of the 41 toxins present in at least half of *Mantella laevigata* individuals in this study.** Abundance is represented by the integrated area under the curve (no units) from the mass spectrometry ion chromatogram for each toxin and frog.

**S5 Table. Detailed sample information for barcoded ant species.** Both morphologically and genetically identifiable ant specimens with exemplar photos representing respective taxonomic groups across seasonal groups are listed with their sample ID’s.

**S6 Table. Detailed sample information for barcoded arachnid species.** Genetically identifiable arachnid specimens with exemplar photos representing respective taxonomic groups across seasonal groups are listed with their sample ID’s.

**S7 Table. Detailed sample information for barcoded “other” prey items.** Genetically identifiable ant specimens with exemplar photos representing respective taxonomic groups across seasonal groups are listed with their sample ID’s.

**S8 Table. Detailed literature information on toxins.** For each of the 41 identified toxins, we list the proposed arthropod origin and whether the toxin has been previously found in other mantellid species or other anurans.

